# Benchmarking enrichment and depletion methods for quantitative plasma proteomics in different plasma types and the correlation to clinical routine assays

**DOI:** 10.1101/2025.11.03.686186

**Authors:** Anders Handrup Kverneland, Ole Østergaard, Luisa Marie Schmidt, Mia Østergaard Johansen, Steffen Ullitz Thorsen, Ruth Frikke Schmidt, Christina Christoffersen, Jesper Velgaard Olsen

## Abstract

Plasma proteomics based on mass spectrometry has great potential for biomarker discovery. Plasma is challenging for mass spectrometry due to high dynamic range in protein abundance. Several workflows have been developed to overcome this, and in this study, we compare prominent workflows using platelet-poor-(PPP), platelet-rich plasma (PRP) and serum (SER).

Our results show that depletion workflows including Top14 depletion and acid precipitation allow quantification of very different proteomes than methods based on enrichments of extracellular vesicles such as bead-based enrichment or ultracentrifugation. Enrichment methods are superior in terms of proteome depth and quantitative performance but may be less robust in large cohorts. There is a very high correlation between PPP and PRP samples with all methods and less to SER samples - especially with enrichment workflows. The correlation of 10 protein measurements, performed by clinical routine processes on a Cobas system, showed heterogeneous results. Low abundant proteins with biological dynamics within a healthy cohort, including c-reactive protein and lipoprotein(a), correlated very well to proteomic workflows while others, including albumin and transferrin, correlated poorly.

In conclusion, the workflow for plasma proteomics should be aligned with the aim of the analysis and the setup of the sample collection.

Graphical Abstract

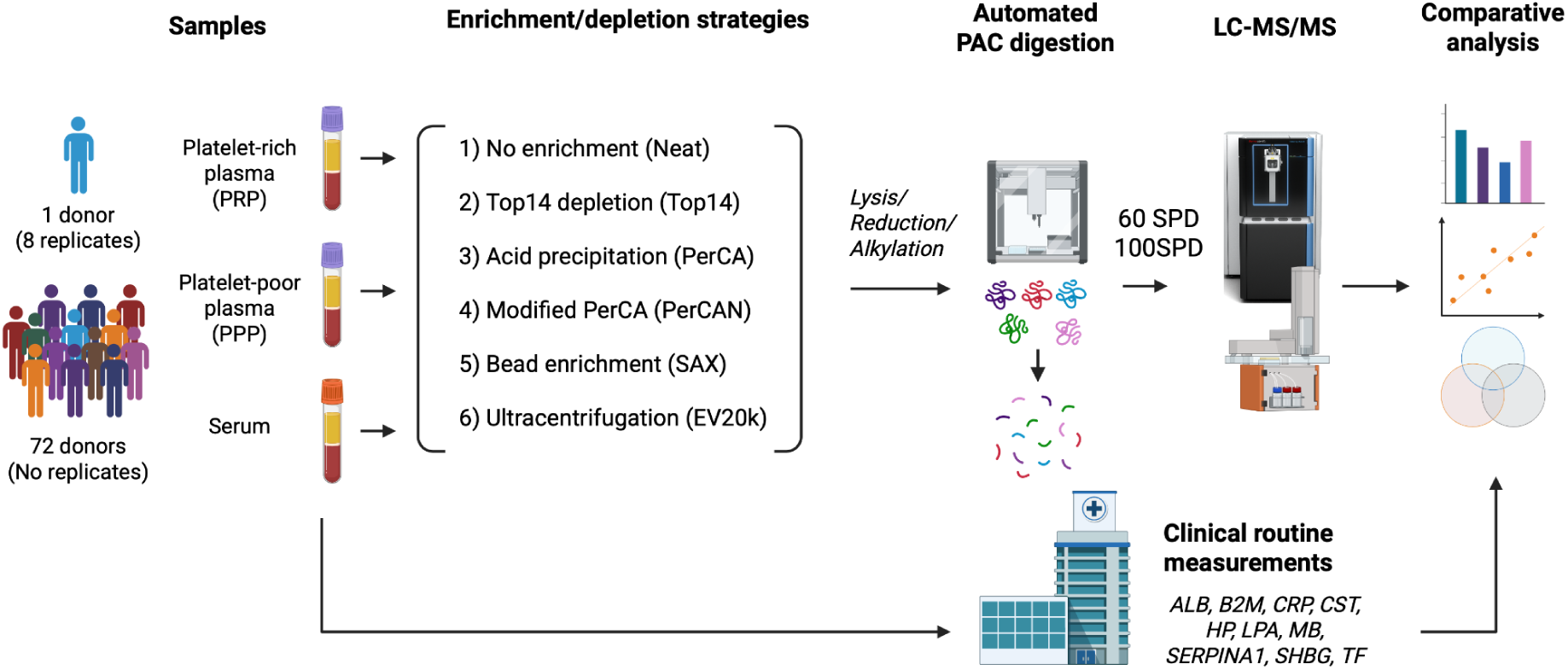

## Background

Peripheral blood is the single most important body fluid for measuring biomarkers in clinical practice. It is a rich source of systemic biological information and is firmly implemented into daily clinical practice given its relative easy accessibility. For measurements of proteins and other molecules, the non-cellular component of the blood is isolated through centrifugation into either serum or plasma. Plasma is obtained by centrifugation after blood has been collected in a tube containing an anticoagulant such as EDTA or citrate to prevent clotting. Conversely, serum is the fluid that remains after a blood sample has been allowed to clot, and cells and resulting fibrin clot are removed by centrifugation and thus contains the same constituents as plasma but devoid of most coagulation factors. Both plasma and serum are attractive biospecimens for novel biomarker discovery.

Plasma proteins have important physiological functions including coagulation, signaling, immunity and the combined concentration of blood proteins is important for maintaining blood osmolarity. In clinical practice, protein-based biomarkers are measured in the clinical biochemistry department. Quantitative assays are usually highly automated and based on detection of single analytes using specific immuno-chemical reactions followed by measuring light scattering as with the Cobas system ^1^.

In parallel, proteomic methods for simultaneous measurement for 100s or even 1000s of proteins are in rapid development. Notable methodologies include affinity-based commercial platforms such as SomaLogic and Olink, where aptamers or antibodies specific for a single protein target are combined into large screening panels and quantified using next generation sequencing (NGS) techniques^2,3^.

Proteomic measurements based on liquid chromatography coupled tandem mass spectrometry (LC-MS/MS) represent a different approach to proteome measurements and offer an unbiased approach to protein identification and quantification. In LC-MS/MS, peptides are separated by reversed-phase chromatography and continuous elution using a gradient increasing in organic solvent before ionization by electrospray. The peptide ions are measured by their mass-to-charge (m/z) ratio and identified by their fragmentation pattern in the mass spectrometer. The most common workflow is bottom-up proteomics, where the proteins are cleaved into peptides by tryptic digestion before analysis by LC-MS/MS. Label-free peptide quantitation is based on the MS signal intensity measured in the mass spectrometer. The quantified peptides are converted into protein quantities *in silico* using a species-specific protein sequence database containing all expected proteins from the organism studied ^4^.

An advantage of MS-based proteomics over affinity-based platforms is the unbiased measurement of sample constituents without the need for an antibody panel or other predefined target selection. It is not prone to specificity or sensitivity issues seen with affinity platforms. Instead, LC-MS/MS is mainly limited by the dynamic range of the proteins in the samples as well as the acquisition speed of the mass spectrometer and the chromatographic performance of the LC system.

Plasma (and serum) is particularly challenging for LC-MS/MS analysis due to the extreme ranges of the protein abundances. The continuous improvement of both mass spectrometers and LC systems has significantly increased the performance of LC-MS/MS in plasma, but still the achievable proteome depth remains a fraction of what is possible from cultured cell lines and tissue - even at a single cell level ^5,6^, which is a result of MS vendors focussing more on instrument sensitivity than on instrument dynamic range.

Still, MS-based plasma proteomics is an increasingly attractive method for biomarker studies. To mitigate the dynamic range issue in plasma, sample enrichment or depletion workflows are continuously being developed. Depletion methods include depletion of highly abundant proteins using affinity-based methods or by protein precipitation, while enrichment methods increase the concentrations of subcellular particles such as vesicles, platelets or cellular debris using high-speed centrifugation or beads ^7–11^.

In the present study, we assessed and compared the performance of popular MS-based plasma proteomics sample preparation workflows. We specifically wanted to investigate how pre-analytical handling including plasma/serum isolation and different enrichment/depletion techniques impacts the quantifiable plasma proteome. To accurately benchmark the different methods against each other, we quantified the intra-sample variation across multiple methods, and assessed the inter-individual performance of the different methods across a healthy donor cohort. For an external and clinically relatable reference and for method evaluation, we performed individual measurements of 10 selected proteins in the same samples at the hospital laboratory using clinical routine workflows.

## Method

### Sample collection

Samples were collected from anonymous healthy adult blood donors from the Capital Region Blood Bank at the Copenhagen University Hospital-Rigshospitalet. Plasma was collected in 4 mL K2-EDTA tubes (Greiner Bio-One) and centrifuged for 6 min at 2000g as platelet-rich plasma (PRP) or at 3000g as platelet-poor plasma (PPP). Serum (SER) samples were collected in 4 mL dry tubes and left to coagulate for 30 min before centrifugation for 6 min at 2000g. The 6 min centrifugation time corresponded to 10 min centrifugation in 2.5 cm longer 6 or 9 mL tubes. All samples were processed within 2 hours after collection, aliquoted and frozen at −80 °C until further analysis.

### Sample processing workflows

The donor samples were processed either as neat plasma/serum (Neat) by different enrichment or depletion workflows recently described in the literature for plasma proteomics. For the depletion workflows resin-based depletion of highly-abundant proteins (Top14) or acid precipitation using perchloric acid in the original version (PerCA) or the modified version (PerCAN) was used, while the enrichment workflows encompassed extracellular vesicle enrichment by magnetic strong-anion exchange beads (SAX) or by ultracentrifugation at 20.000g (EV20k) ^7,8,12,13^

Neat samples were denatured, reduced and alkylated in sample buffer (4% SDS, 100 mM TrisHCl pH 8.0 supplemented with 5 mM TCEP and 10 mM CAA; 1% SDS final concentration) for 15 min at 37 °C The samples were then diluted 250X (final concentration) with PBS (Gibco, PBS pH 7.2).

For the Top14 depletion, 5 uL sample were added to 200 uL Depletion Resin (High-Select™ Top14 Abundant Protein Depletion Resin, Cat# A36372, Rockford, IL) in 96-well filter plates (Millipore cat. no. MSHVN45) and incubated 10 min at ambient temperature with frequent mixing. The samples were then centrifugated for 2 min at 800g, and 30 uL sample was transferred from the high-abundance protein depleted filtrate to a 96-well plate followed by addition of 90 uL sample buffer for reduction and alkylation.

For acid precipitation (*PerCA*), 50 uL sample was mixed with 450 uL Milli-Qfollowed by addition of 25 uL 70% Perchloric Acid (PerCA; final concentration 3.5%) before incubation at −20 °C for 15 min. The samples were then centrifuged at 3200g for 60 min at 4 °C and 250 uL from the supernatant was mixed with 40 uL 1% TFA and loaded on a uSPE HLB plate (Waters, Cat# 186001828BA) preconditioned with 100 % methanol and 0.1% TFA. After washing 3 times with 0.1 % TFA, the proteins were eluted with 90% ACN supplemented 0.1%TFA followed by vacuum centrifugation until dryness. Finally, the dried pellets were resuspended in 35 uL 50 mM ammonium bicarbonate for digestion with trypsin and eLysC (As described by^8^).

In addition to the PerCA workflow, we also performed the recently modified PerCA workflow that includes a pH neutralization step using NaOH (PerCAN)^12^. In brief, 10 uL plasma was diluted with 40 uL Milli-Q water in a 96-well plate before addition of 50 uL 1.0 M PerCA and incubation at 4 °C for 60 min. The plate was then centrifuged for 60 min at 4,000g in a cooled centrifuge (4 °C) followed by transfer of 48 uL supernatant to a new plate and addition of 16 uL 1.4 M NaOH titrated to neutralize the supplemented PerCA. Subsequently, reduction and alkylation were performed by addition of a tetra-ethyl ammonium bicarbonate (TEAB) based buffer (60 mM) supplemented with TCEP (5 mM) and CAA (10 mM, all final concentrations) and incubation at 95 °C for 10 min before cooling and addition of trypsin and eLysC.

For the bead enrichment (SAX), 20 uL or 40 uL sample was diluted 1:2 with binding buffer (100 mM Bis-Tris Propane, pH 6.3, 150 mM NaCl) in a 96-well plate. Then 100 ug (5 uL) SAX beads (ReSyn Bioscience, Cape Town, South Africa) suspended in binding buffer, were added to each sample and incubated for 60 min on a thermoshaker at 600 rpm at ambient temperature. The supernatant was removed using a magnetic rack and the beads were washed three times in a washing buffer (50 mM Bis-Tris Propane, pH 6.5, 150 mM NaCl). The beads were then resuspended in 50 uL sample buffer and incubated for 15 min at 37 °C to complete reduction and alkylation.

For the EV20k enrichment, 500 uL sample was transferred to 1.7 mL microcentrifuge tubes supplemented with 500 uL PBS. To remove remaining platelet or coagulates, the samples were centrifuged at 3000g for 2.5 min and the supernatant was transferred to a new tube. For the EV enrichment, the samples were then centrifuged at 20,000 g for 30 min followed by discarding the supernatant except for last 20 uL above the pellet. After supplementing with 980 uL PBS, the centrifugation was repeated as a single washing step with PBS leaving a ∼20 uL *EV20k* pellet. The pellets were incubated with sample buffer (4% SDS, 100 mM TrisHCl pH 8.0 supplemented with 5 mM TCEP and 10 mM CAA; 1% SDS final concentration) for 15 min at 37 °C before overnight digestion with trypsin and eLysC.

### BCA analysis

Bicinchoninic Acid (BCA) analyses were performed after each preparatory sample workflow to assess the protein yield after the enrichment or depletion steps. The Thermo Scientific Pierce™ BCA Protein Assay Kits (Cat #23225) was used for analysis according to the manufacturer’s instructions and the resulting absorbances were recorded at 550 nM using a FLUOstar® Omega spectrophotometer.

### Sample digestion and EvoTip loading

With exception of the PerCA sample, the samples subject to the neat, enrichment or depletion workflows were digested using protein aggregation capture (PAC) protocol ^14^, desalted and loaded on stagetips (EvoTips, Evosep Biosciences, Odense Denmark) using an OT2 Opentrons Robot as described elsewhere^5^. The scripts for the digestion and stagetip loading were generated using an online script generator tool (https://www.evosep.com/support/automation-opentrons-ot2/). For the automated workflow, 5 uL sample was diluted to approximately 0.5 ug/uL, transferred to a sample plate together with 100 ng Hydroxyl-beads (ReSyn Biosciences) and placed on the OT2 robot. The script was set to load approximately 500 ng sample peptide on the stagetip. For the SAX samples, 10 uL sample (including 20 ng SAX beads) were used without Hydroxyl-Beads. The PerCA and PerCAN samples were digested in 50 mM ABC or in 60 mM TEAB buffer and manually loaded onto EvoTips.

### LC-MS/MS analysis

The peptide samples were eluted from Evotips using an Evosep One LC system (Evosep Biosystems) and separated using an Evosep 8 cm (EV1109, Evosep) performance column connected to a steel emitter (EV1086, Evosep) and heated to 40 °C over an 11 min (100 SPD) or 21 min (60 SPD) gradient.

The eluted peptides were electrosprayed into an Orbitrap Astral mass spectrometer (Thermo Fisher Scientific) using 1800 V spray voltage, funnel radio frequency level at 40, and a heated capillary temperature set to 275 °C in positive mode. Full scan precursor spectra (380–980 Da) were recorded in profile mode using a resolution of 240,000 at 200 m/z, a normalized automatic gain control (AGC) target of 500%, and a maximum injection time of 3 ms. The fragment spectra were acquired in narrow-window data-independent acquisition (nDIA) mode, with a precursor mass range of 380 to 980 m/z with 2 Th isolation windows without overlap, in total 299 scan events per scan cycle ^15^. Isolated precursors were fragmented in the HCD cell using 25% normalized collision energy, a normalized AGC target of 500%, and a maximum injection time of 2.5 ms.

### Clinical biochemistry analysis

Protein quantifications at the hospital clinical biochemistry department were performed using standardized and validated assays for clinical use at the Department of Biochemistry *(Clinical routine)*. Ten proteins were measured including: Albumin (ALB), alpha-1-antitrypsin (SERPINA1), high sensitive C-reactive protein (hs-CRP or simply CRP), Cystatin-C (CST3), Haptoglobin (HP), lipoprotein(a) (LPA), Beta-2-microglobilin (B2M), Myoglobin (MB), Transferrin (TF) and sex-hormone binding globulin (SHBG). Measurements were performed using Cobas® Pro c502, c503, and e801 systems from Roche Diagnostics. ALB was measured by pH-dependent colorimetry; SERPINA1, HP, B2M, and TF were measured using immunoturbidimetry; CRP, CST3, LPA, and MB were measured using particle-enhanced immunoturbidimetry, and SHBG were measured by a sandwich-based electrochemiluminescence immunoassay. The proteins were quantified using reagent kits from Roche with the following product codes: ALB: 08056714190; SERPINA1: 08101396190; CRP: 08057605190; CST3: 08105596190; HP: 08106045190; LPA: 08106126190; B2M: 08105529190; MB: 07027583190*; TF: 08058733190; and SHBG: 07258496190.

### Data analysis

The MS RAW-files were analyzed using DIANN v2.2.0 (academia) searching against a FASTA file containing the human proteome (SwissProt, 20,421 sequences, downloaded on May 6th 2025) including two amino acid sequences from the applied proteolytic enzymes ^16^. Default search settings were used. In brief, carbamidomethyl was set as a fixed modification and *N*-terminal acetylation and oxidation of methionine as variable modifications. Trypsin was specified as the proteolytic enzyme with a maximum of one missed cleavage allowed. The mass tolerance was set to automatic inference at both MS1 and MS2 levels with a precursor FDR level at 1%.. Quantification was performed using the Quant UMS (high precision) setting. Intensity-Based Absolute Quantification (IBAQ) was calculated after the DIANN search by dividing the label-free quantification by the number of theoretical peptides for each protein group.

The data analysis was performed in R version 4.2.2 (R Core Team (2021), R: A language and environment for statistical computing, R foundation for statistical computing) with R Studio 2023.09.1 Build 494. The patient data was normalized using the variance stabilization normalization using the vsn package ^17^. The statistical analysis was performed using the rstatix package. Gene Ontology enrichment analyses were performed with the clusterProfiler package ^18^. Subcellular locations were predicted using the DeepLoc prediction tool ^19^. Protein annotations were.

### Experimental Design and Statistical Rationale

To obtain reliable performance estimates – evaluated as the number of quantified peptides and proteins – and for estimating the uncertainty of the protein quantifications – evaluated as the %CV on the quantitative readouts – we evaluated all workflows by performing 8 workflow replicates using aliquots from the same QC specimen (either PPP, PRP and SER) as starting material.

To gain further insight into the performance of the methods evaluated in the study and to obtain insight into the variation of protein abundances in a medium sized healthy cohort, we analyzed 72 donor samples (36 PPP, 36 PRP and 72 SER samples) using the 6 evaluated workflows with one workflow replicate per sample. The size of the cohort was primarily limited by the availability of human samples for processing but we found the number sufficient for the current analysis.

## Results

### Proteome depth and correlation between methods

To assess the achievable depth of the plasma proteome in 21 min LC-MS/MS measurement time and evaluate the reproducibility of the different sample preparation methods, we analyzed a PRP sample in 8 workflow replicates using aliquots derived from the same starting material from one healthy donor. The proteome depth ranged from 724 proteins in neat plasma to 4944 proteins in EV20k plasma (Figure 1A, median values). All workflows were able to identify a high number of FDA drug targets across the whole intensity range, and the number increased gradually with increasing proteome depth (Figure 2B). We compared the quantitative performance by assessing the proportion of proteins with low to high coefficients of variation (CV). Neat plasma had the highest proportion of proteins with low CVs (Figure 3C). In all methods, the majority of proteins had CVs below 20% with the exception of PerCAN where the CVs where the CVs were generally higher.

**Figure 1:**
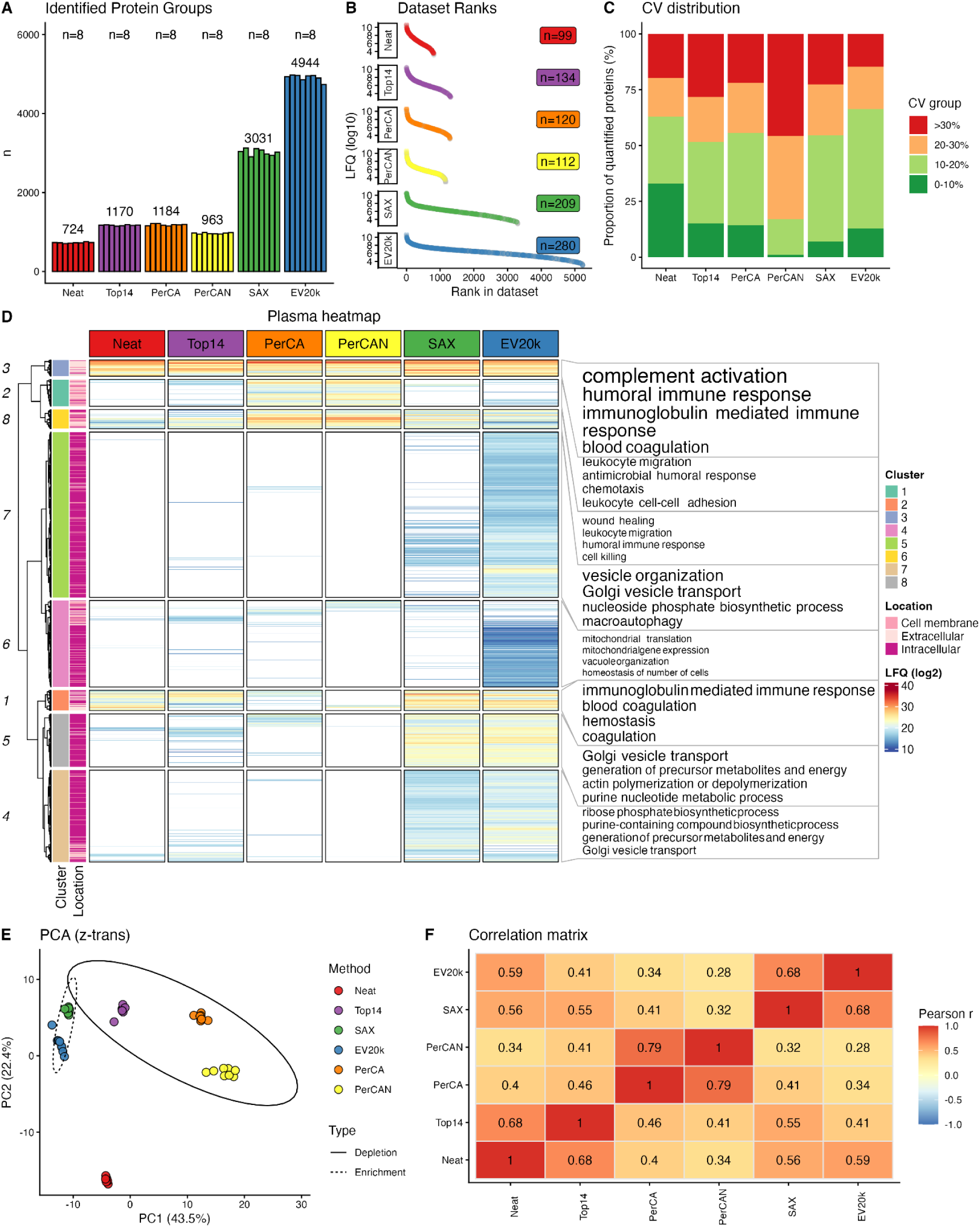
Overall comparison of enrichment and depletion methods. A: Quantified Protein IDs in the plasma proteome in workflow replicate samples (n=8) that were run in parallel using the different methods of enrichment and depletion. The median ID count is shown. B: Label-free-intensity vs data set rank of identified protein groups. The FDA targeted proteins are colored and the number is shown in the label for each workflow. C: Grouped distribution of Coefficients of Variation (CVs) within each method. D: Heatmap and clustering of the protein identified in all replicates within each method. The top 3 Enriched GO terms are shown on the right to each cluster with text size relative to decreasing q-value (interval: 3e-48 to 4e-9). Missing values are shown in white. For clustering, the missing values were imputed using left-shifted Gaussian imputation. Protein Groups were hierarchically clustered using Euclidean distance and Ward’s minimum variance method. E: Principal Component Analysis (PCA) of the shared proteins in the plasma proteomes after the enrichment/depletion using the described methods after z-score transformation. F: Correlation matrix based on median intensity of shared proteins between each method. The Pearson’s correlation coefficient is shown in each square.

**Figure 2:**
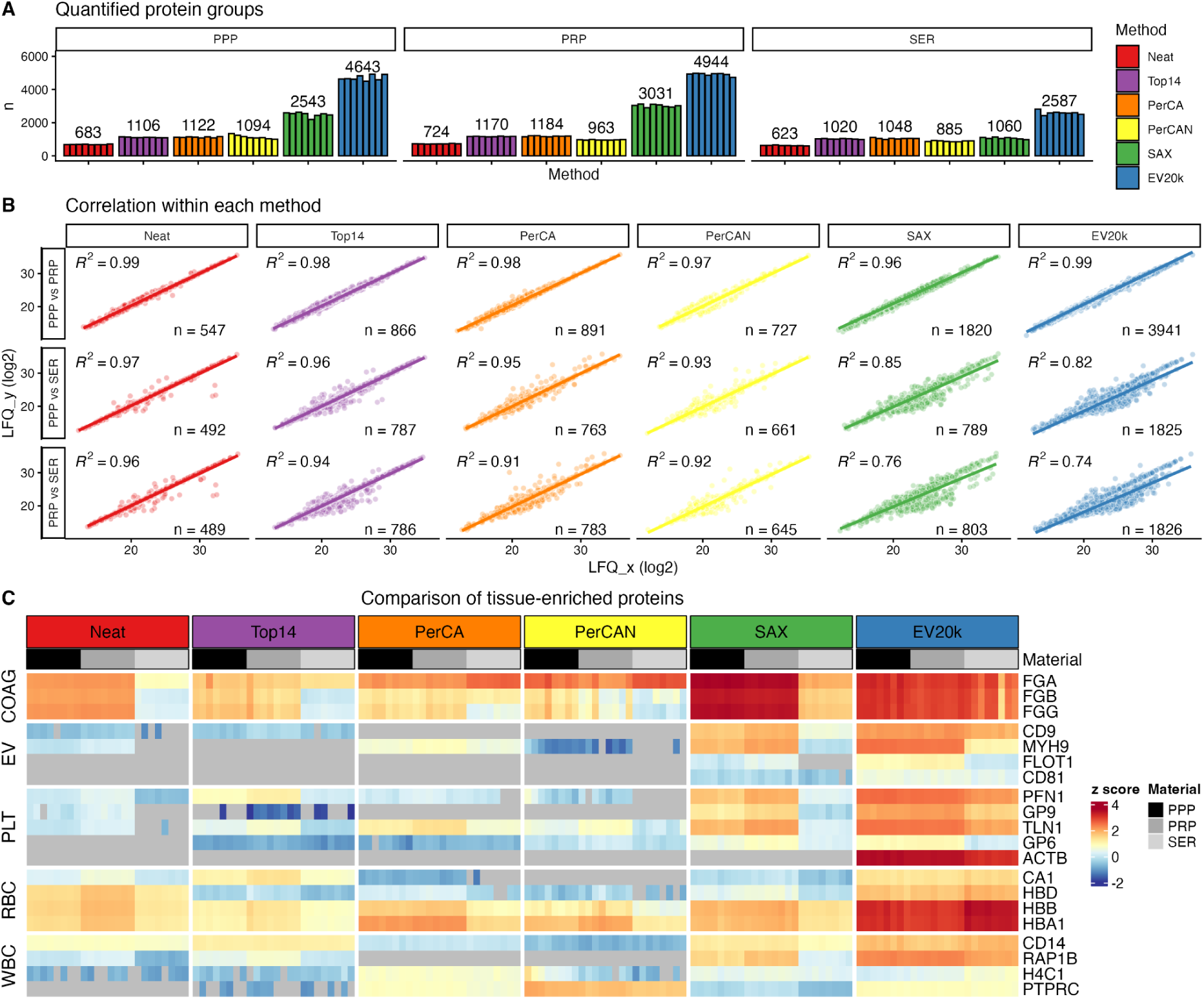
Impact of material type on the quantified proteome after workflow processing. A: Number of quantified protein groups with each method from different material types in 8 workflow replicates processed in parallel. The median counts of quantified proteins are shown. B: Correlation of shared proteins between different material types within each method. The square Pearson correlation coefficient is shown with the number of proteins that was compared. C: Scaled label-free quantification (z-score) intensity of annotated tissue-enriched proteins within each method and each material type. Grey fields represent missing values. PPP: Platelet-poor plasma, PRP: Platelet-rich plasma, SER: Serum, COAG: Coagulation, EV: Extracellular vesicles, PLT: Platelets, RBC: Red blood cells, WBC: White blood cells.

**Figure 3:**
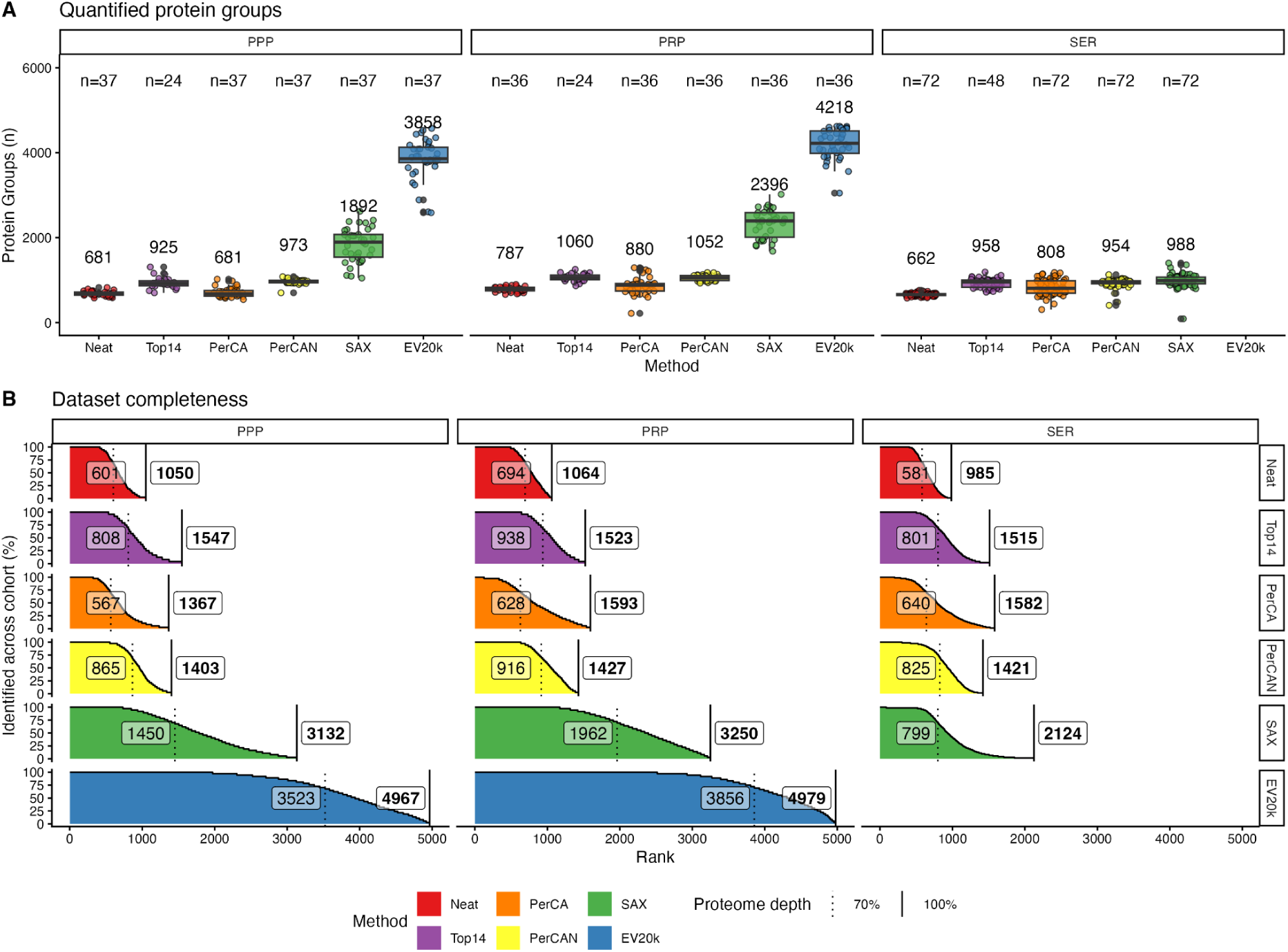
Plasma proteomes in a donor cohort after enrichment and depletion based workflows. A: Number of quantified protein groups after sample processing using enrichment or depletion based workflows stratified on different starting materials. The median protein count for each workflow is shown together with the number of donor samples (n). B: Visualisation of the dataset completeness obtained for each method. The absolute number of proteins found in 70% of samples is shown at the dotted line. The total number of unique protein groups identified in the dataset is shown in bold. PPP: Platelet-poor plasma, PRP: Platelet-rich plasma, SER: Serum.

To find similarities and differences in the Protein Group identifications across the different methods, we performed an unsupervised hierarchical clustering of protein intensities and visualized all obtained plasma proteomes in a heatmap (Figure 3D). This analysis grouped proteins into eight main clusters and by performing a gene ontology (GO) enrichment analysis for each cluster, we found that the protein clusters 3 was shared between all methods and consisted of highly abundant proteins related to complement and immunoglobulins. Notably, one cluster (Cluster 2) enriched in coagulation-related proteins was less abundant in the PerCA and PerCAN workflows, whereas these workflows specifically enriched leucocyte-related proteins (Cluster 1). The EV20k and SAX enrichment workflows shared clusters associated with vesicle transport and mitochondria (Cluster 5), while the EV20k had the addition of endosomal-associated pathways (Cluster 4).

When comparing the obtained proteomes by subjecting the shared proteins to a principal component analysis (PCA), we saw that neat plasma separated itself in principal component 2 (PC2) while PC1 separated the enriched from the depleted plasma (Figure 1E). In PC1, accounting for 41.3% of the variance, the Top14 depletion was very similar to neat plasma. In a pairwise correlation analysis of the median protein intensities, neat plasma correlated strongest with Top14 depleted plasma, while PerCA correlated poorly with a Pearson r < 0.5 to the other methods (Figure 1F). We saw a strong correlation between the PerCA and PerCAN methods and between the enrichment methods.

### Impact of starting material

The pre-analytical sample processing affects the relative content of specific proteins. We wanted to assess the difference between plasma, rich (PRP) or poor (PPP) in platelets, and serum (SER) from the same healthy donor.

The deepest plasma proteome was generally obtained from PRP samples across the different sample preparation methods with close to five thousand proteins identified in EV20k (Figure 2A). We also found that the Neat samples showed very high correlation between all starting materials, with PRP vs PPP having a 0.99 correlation coefficient (Figure 2B). The depletion methods (Top14 and PerCA) showed similar correlation between all different starting materials while the enrichment methods (SAX and EV20k) had a very high correlation between PPP and PRP samples, but relatively poor correlation to SER samples.

Particularly the content of platelets (PLT), extracellular vesicles (EVs), coagulation factors (COAG), red blood cells (RBC) and white blood cells (WBC) is affected by sample processing. We evaluated this, by comparing the relative quantification of commonly associated proteins (Figure 2C). As expected, coagulation factors were depleted in SER samples for all workflows with the exception of the coagulation factor FGA in the PerCA depletion workflow. The PerCAN plasma- and all SAX samples were very high in fibrinogens. Reassuringly, the comparison confirmed that EV proteins were highly enriched in both the EV20k and SAX workflows. As expected, PLT proteins were less abundant in PPP samples compared to the PRP and SER samples and particularly high after EV20k and SAX enrichment.

A PCA together with comparison of CV, subcellular location and plasma starting volume for SAX enrichment can be seen in Supplementary figure 2.

### Performance in a healthy cohort

A major challenge in MS-based plasma proteomics is the considerable inter-individual variance. We tested the performance of the assessed workflows and influence of starting material within a mid-sized human cohort of healthy blood donors (n=72). The age and sex distribution can be seen in Supplementary Figure 3 Due to limited sample material, the EV20k enrichment was not performed in the SER samples and there was only one donor with both PRP and PPP samples for comparison with SER. For the rest of the samples across the cohort plasma samples (either as PRP or PPP) were available for comparison to its matched serum sample.

First, we examined whether the depth of the quantifiable proteome was on par with the proteomes obtained with the workflow replicates presented above. When comparing the median number of protein IDs, we found that the proteome depth was comparable, but slightly lower in the donor cohort (Figure 3A). We observed the highest variance in protein depth within the EV20k PPP samples.

Another common issue for statistical comparisons in plasma proteomics datasets is missing values. To test how variably the different methods could identify and quantify proteins across our healthy sample cohort, we compared the dataset completeness in terms of quantified protein intensity values. Dataset completeness was defined as the number of protein groups quantified in at least 70% of the samples - commonly used as cut-off. With most methods, the relative number of proteins identified in 70% of samples for each method were around 40-60% of the total number of proteins identified for that method (Figure 3B). Surprisingly, it was 70-80% for the EV20k samples - despite having the highest number of protein IDs. PRP showed the overall highest dataset completeness while PPP and SER were on comparable levels except for SAX that showed much a lower level in SER.

### Correlation to clinical biochemistry

To assess the relevance of the quantitative values of the proteins identified with the different sample preparation methods, we compared our MS-based protein measurements to validated clinical routine assays performed at the hospital laboratory. Thus, we identified ten proteins used in clinical diagnostics that are often quantified in MS plasma proteomic datasets and spanning a wide range of physiological concentrations.

The hospital protein measurements of the cohort SER samples (n=72) and their reference levels using the Cobas modular clinical chemistry and immunochemistry analyzer systems can be seen in Supplementary Figure 4A. The ten selected proteins could be quantified in most, but not all, samples with both routine diagnostics and mass spectrometry. Myoglobin (MB) was missing in all SAX samples and more than half of the EV20k samples. Apolipoprotein-A (LPA) values were missing in more than a quarter of the hospital measurements. The low abundant proteins, C-reactive protein (CRP), Cystatin-C (CST3), Sex hormone-binding globulin (SHBG) and MB, were missing in many PerCA samples. The fraction of missing values for each method is shown in Supplementary Figure 4B.

The proteins spanned approximately six orders of magnitude in dynamic range in protein concentrations in serum (Figure 4A). The ten proteins were ranked based on their median abundance quantified by the clinical routine assay and compared to the corresponding abundance ranks in PRP determined by mass spectrometry (Figure 4A and Figure 4B) after processing by the six assessed workflows. We found that the ranks and magnitude of dynamic range were largely retained from the hospital assay to the MS data.

**Figure 4:**
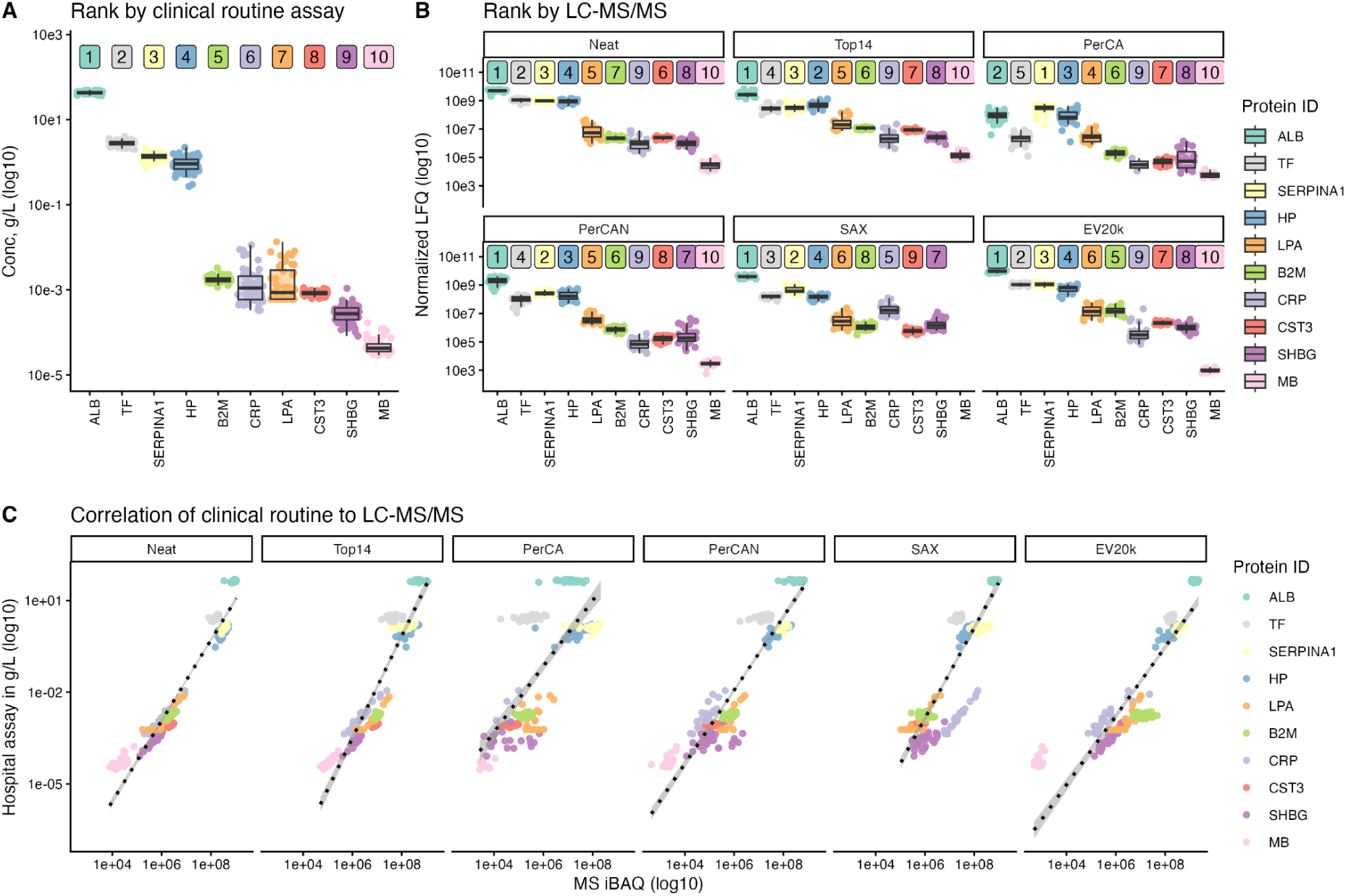
Overall comparison of mass spectrometry based estimates and clinical routine measurements for 10 selected proteins. A: Boxplot of protein concentration (ug/uL) measured by clinical routine analysis in SER samples. The relative rank is shown in the colored label. B: Boxplots of protein LFQ in PRP samples after processing by the indicated workflows prior to MS analysis. The relative rank of the protein is shown in the colored label. C: Correlation of the absolute concentrations measured by clinical routine assays in serum samples and the Intensity-Based Absolute Quantification (iBAQ) measured by mass spectrometry after processing PRP by each of the evaluated enrichment or depletion workflows. The dashed line is based on linear regression. LFQ: Label-free Quantification.

Albumin (ALB) was the most abundant protein by all methods except PerCA. CRP was consequently less abundant in the MS results than by the hospital assay, which might be attributed to the high fraction of limit-of-detection (LOD) values in the hospital assay.

Next, we correlated the MS-based protein abundances, estimated using intensity-based absolute quantification (iBAQ), to the clinical routine assay concentration measurements after logarithmic transformation (Figure 4C). We found a linear correlation with high correlation coefficients for all methods. The best correlation between MS-based protein abundances and clinical routine assays was found between clinical routine values in serum and LC-MS/MS analysis of neat plasma (R^2 = 0.94) while the poorest correlation was with the PerCA values (R^2=0.58).

In figure 4C, specific proteins such as ALB appeared poorly correlated between the hospital assay and each of the MS methods. We investigated this further by visualizing the correlatings between the results for each protein individually. A heatmap of all single protein correlation coefficients between clinical routine values and LC-MS/MS-based estimates is shown in Figure 5A. The results illustrate that the correlations were highly protein analyte dependent. As examples, the correlations of clinical routine with LC-MS/MS estimates of LPA, CRP and SHBG are shown in Figure 5B. LPA is highly correlated to all methods except PerCA although many clinical routine measurements were missing and presumably below the limit of detection. Clinical CRP values correlated well with LC-MS/MS values from each workflow - although the PerCA workflow only resulted in CRP estimates from 3 samples. For SHBG, clinical routine estimates correlated well to LC-MS/MS values after the NEAT, Top14 and EV20k workflows, but not with values from the PerCA, PerCAN or SAX workflows. The correlation plots for all ten proteins is shown in Supplementary Figure 5.

**Figure 5:**
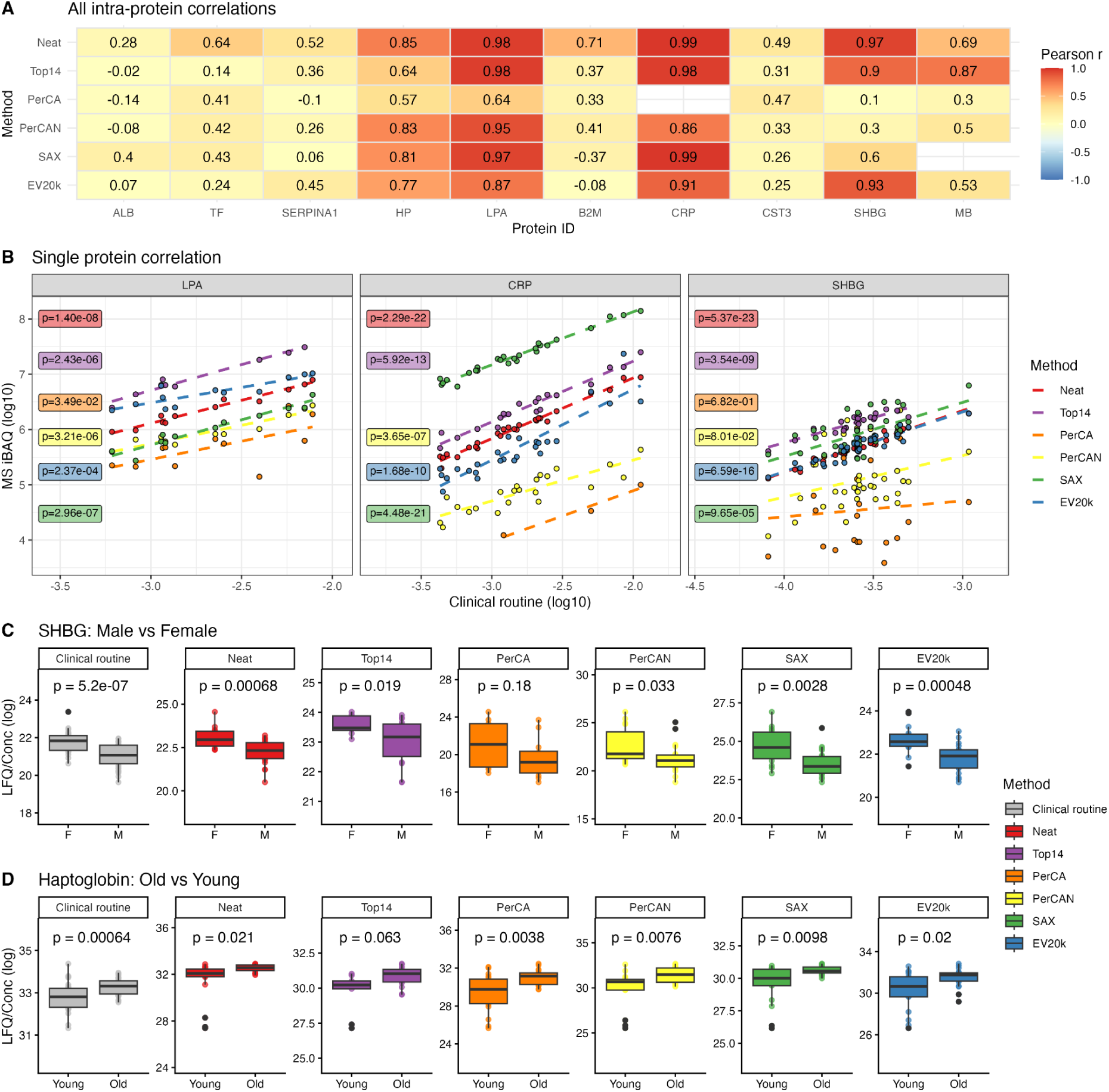
Correlations of abundance levels for ten selected proteins between levels quantified by clinical routine analysis and values obtained after LC-MS/MS analysis in combination with six evaluated workflows. A: Correlation between the hospital measurements and MS processing workflows for LPA, CRP and SHBG. B: Visualisation of the correlations of the absolute concentrations measured by the hospital laboratory with the Intensity-Based Absolute Quantification (IBAQ) values measured by mass spectrometry for each measured protein and each enrichment or depletion method. The square Pearson r correlation coefficient is shown. C: Boxplots showing Sex Hormone Binding Globulin (SHBG) levels measured by the hospital assay and by mass spectrometry stratified by sex. The p-values are calculated by uncorrected student’s t-test. D: Boxplot of Haptoglobin (HP) levels measured by hospital assay and by mass spectrometry stratified by sex. The p-values are calculated by uncorrected student’s t-test.

We next looked into how donor characteristics such as age and gender would be reflected into the levels of the selected proteins quantified by the clinical routine method and by LC-MS/MS after processing the samples by the six assessed workflows. The SHBG protein abundance was highly reflected by sex, and when comparing male to female values estimated by clinical routine measurements, we could see significantly higher levels in females (Figure 5C). Similarly, we observed statistically significant different SHBG estimates between males and females by LC-MS/MS analysis after sample processing by the assessed workflows, except for the PerCA workflow. Similarly, we tested the age effect on Haptoglobin (HP) levels by comparing the old vs young donors divided into two groups by the age median (Figure 5D). In this analysis, only the Top14 workflow, where HP is selectively depleted, were unable to reproduce the significant difference between the age groups observed with the clinical routine measurements. All effects of sex and age on estimated protein levels determined by clinical routine measurements is shown in Supplementary Figure 6.

## Discussion

MS-based plasma proteomics holds a huge potential for clinical applications and biomarker discovery. As demonstrated here and by others, the challenge of the high dynamic range of protein abundances in neat plasma can be partly overcome through several competing depletion/enrichment steps but with the risk of affecting sample integrity. In this study, we assessed six workflows intended for sample processing prior to LC-MS/MS based analysis, compared them to each other and benchmarked the results with single protein measurements obtained by validated clinical routine analysis using a Cobas system.

In terms of proteome depth, all our tested methods performed at expected or higher levels as reported by others ^7,8,10,11,13,20^. When comparing the different sample preparation workflows before MS analysis, the deepest proteome was achieved through EV enrichment methods without compromising reproducibility based on CV values. This was in contrast to results obtained by other groups, who observed high(er) CVs between workflow replicates for EV isolation. One plausible explanation for this discrepancy, could be the practical laboratory handling of the EV pellets handling after each centrifugation step or the extent supernatant removal.

Unsurprisingly, and across all methods, the highest number of proteins could be identified and quantified from platelet-containing plasma (PRP), where platelet associated proteins contribute to the observed plasma proteome. More surprisingly was the finding that the abundance correlations of shared proteins between different material types (SER, PRP and PPP) were high in both the neat and processed material. The intra-individual correlation between PRP and PPP obtained after enrichment/depletion workflows remained very high and suggests some tolerance to pre-analytical variability during sample handling.

Overall, when focussing solely on protein abundances, the MS-based estimates correlated well with values obtained by clinical routine analysis over the 5 orders of magnitude represented by the 10 selected proteins. However, when looking closer into the correlations between MS based estimates for each of the ten selected proteins individually, only a subset of the ten proteins showed high correlation with the clinical assays. The low physiological dynamics in a healthy cohort for some of the ten selected proteins had the effect that the physiological variance within the cohort was comparable to the workflow variance leading to less obvious correlations. Indeed, the highest correlation between clinical estimates (Cobas system) and the LC-MS/MS based estimates seemed to be for proteins showing sex- and age-dependent concentration variations across the cohort as was the case for SHBG and myoglobulin. Similarly, part of the hospital assays were not optimized for very low concentrations of disease related biomarkers exemplified by the many LPA and CRP measurements below the LOD of the Cobas system.

Our results demonstrate that there are many ways to dig deeper into the plasma proteome and that they can be performed without severely impacting sample integrity. It is also clear that sample processing workflows based on enrichment or depletion have different effects on the resulting proteome and that the purpose of analysis–for example the disease or physiological condition studies–should be considered before deciding the preferred workflow. While enrichment methods give more protein IDs, these are primarily intracellular proteins, and would most likely be relevant to the biomedical questions related to the EVs. In our results, the depletion-based workflows performed lower in terms of quantified proteins and also showed somewhat higher CVs but also had the benefit of showing higher robustness to starting material (serum vs. plasma) than the enrichment methods.

Another important consideration is how well the assessed workflows can cope with increasing sample numbers. The superior proteome depth obtained by isolating extracellular vesicles may seem attractive, but since this workflow in its current form includes a lot of manual labor, it may be unfeasible to perform on large cohorts including thousands of samples if it involves ultracentrifugation.

Here the SAX base workflow seems more attractive as the magnetic bead base format is easier to automate in a 96-well plate based format. Large cohorts often include sample collection at multiple centers over extended time spans, and they have an increased risk of variability in e.g. centrifugation speed, tube sizes and time to sample handling, which leads to increased pre-analytical variability. Our study showed that the depletion based workflows were more tolerable to this kind of variation. This is in agreement with a recent publication that also demonstrated high robustness of the acid precipitation workflow to blood component contamination suggesting that this workflow might be more favorable for multicenter cohorts prone to high degree of pre-analytical variability ^21^. The pros and cons of the 6 assessed workflows are summarized in table 1 for a quick overview.

**Table 1:** Comparison of methods.

|  | Material needed | Cost per sample | Workload per sample | Proteome depth | Quantitative performance | Scalability |
| --- | --- | --- | --- | --- | --- | --- |
| Neat | <1 uL | + | + | + | +++ | +++ |
| Top14 depletion | 5-10 uL | +++ | ++ | ++ | ++ | ++ |
| PerCA | 50 uL | + | + | ++ | + | +++ |
| PerCAN | 5-50 uL | + | + | ++ | + | +++ |
| SAX | 20-40 uL | ++ | ++ | +++ | ++ | +++ |
| EV20k | 500 uL | + | +++ | +++ | ++ | + |

Finally, the overall performance of neat plasma is close to what can be achieved through some laborious workflows in terms of depth and superior in terms of quantitative performance. It is possible that with the continuous technical and computational improvements, the plasma proteome of neat unprocessed plasma might soon be sufficient for many biological applications and studies.

## Acknowledgements

Work at The Novo Nordisk Foundation Center for Protein Research (CPR) is funded in part by donations from the Novo Nordisk Foundation (NNF14CC0001, NNF24SA0098829 and NNF21OC0072070). This project was supported by a center-of-excellence grant from the Danish National Research Foundation to Copenhagen Center for Glycocalyx Research (DNRF196). This project was also supported by a grant from the Danish Agency of Higher Education and Science to establish the PLATO research infrastructure: Danish National Mass Spectrometry Platform for Proteomics and Biomolecular Imaging (5229-00012B).

Ann Kristine Thorsteinsson is acknowledged for her technical expertise. We are grateful for the help from staff at the Blood Bank, unit 9501, who were instrumental for recruiting donors and collecting blood for this project.

## Author contributions

AK, OOE, and LMS prepared all samples and performed proteomics experiments. MVF, ST and CC supervised hospital measurements. AK and OOE analyzed the resulting data and wrote the first version of the manuscript. CC, RFS and JVO critically evaluated the results. All authors read, edited and approved the final version of the manuscript.

## Conflict of interest statement

The authors declare no competing interests.

## Notes

### Competing Interest Statement

The authors have declared no competing interest.

