## Supplementary figures for "Benchmarking enrichment and depletion methods for quantitative plasma proteomics in different plasma types and the correlation to clinical routine assays"

### Supplementary material

#### Supplementary tables

Sup. Table 1: Protein ID and median label-free intensity (LFQ) of all materials across methods and starting materials of the biological replicates

Table1

Sup. Table 2: Top10 GO enrichment after enrichment/depletion workflow in PRP

Table2

Sup. table 3: Protein ID and median label-free intensity in the donor cohort across methods and materials.

Table 3

### Supplementary figures

#### Sup. Figure 1:

A: Upset plot of quantified protein groups and overlap between data sets.
B: Number of quantified protein groups of all methods using 11.5 min (100SPD) and 21 min (60SPD) gradients. The median number of 8 workflow replicates is shown.
C: Number of peptides (stripped sequences) of all methods using 11.5 min (100SPD) and 21 min (60SPD) gradients. The median number of 8 workflow replicates is shown.
D: Denisity plot of the number of precursors per quantified protein groups. The median number is shown.

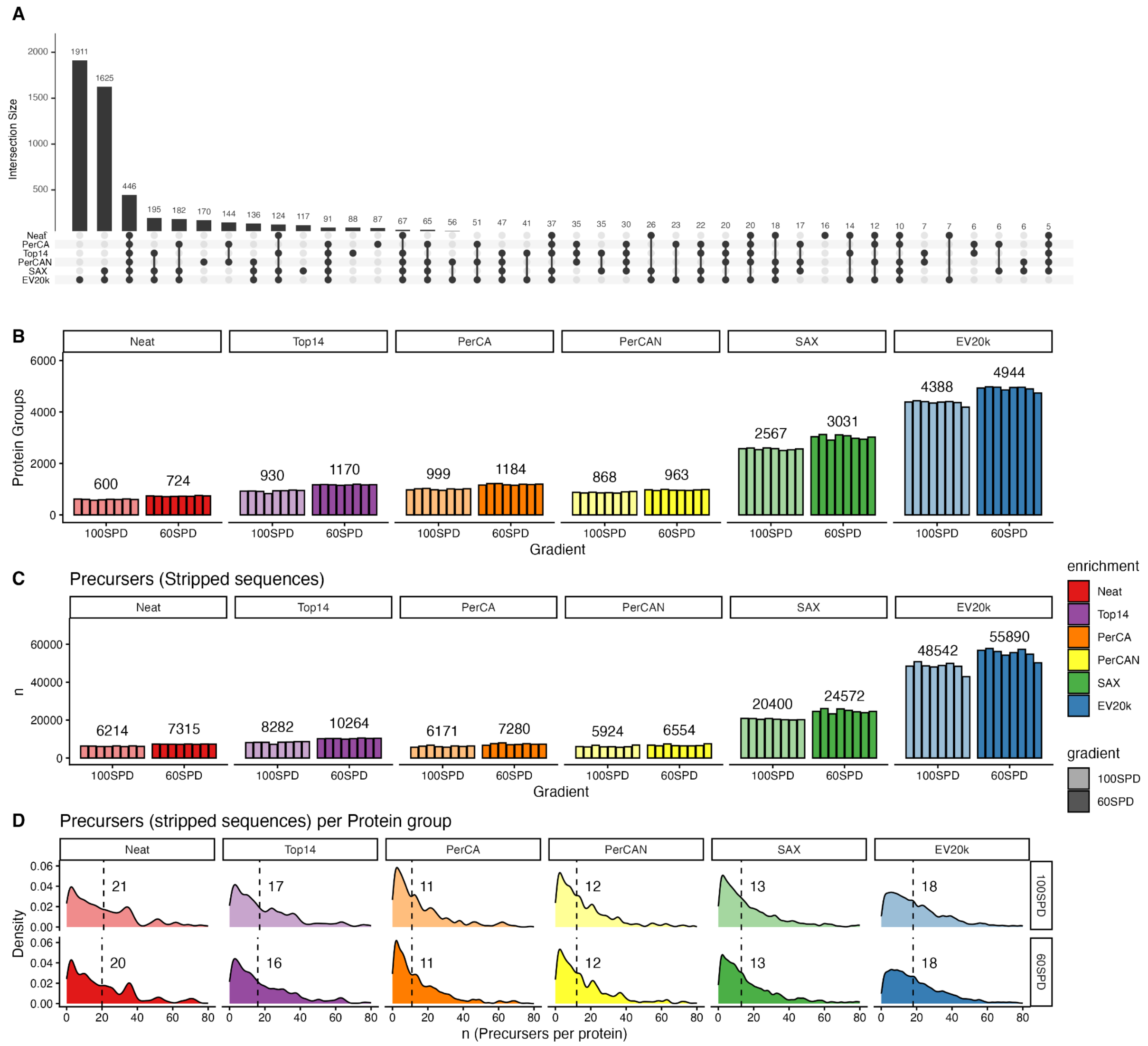

Sup. Figure 2:

A: PC1 and PC2 of a principal component analysis (PCA) of plasma workflows and starting material.
B: PC3 and PC4 of a principal component analysis (PCA) of plasma workflows and starting material.
C: Violin plot of coefficients of variation within each workflow and starting material. The median CV is shown.
D: Stacked boxplot with the distribution of predicted subcellular location within each workflow and start material.
E: Comparison of using 20uL vs 40ul of starting material for the same amount of SAX beads before the SAX enrichment workflow.

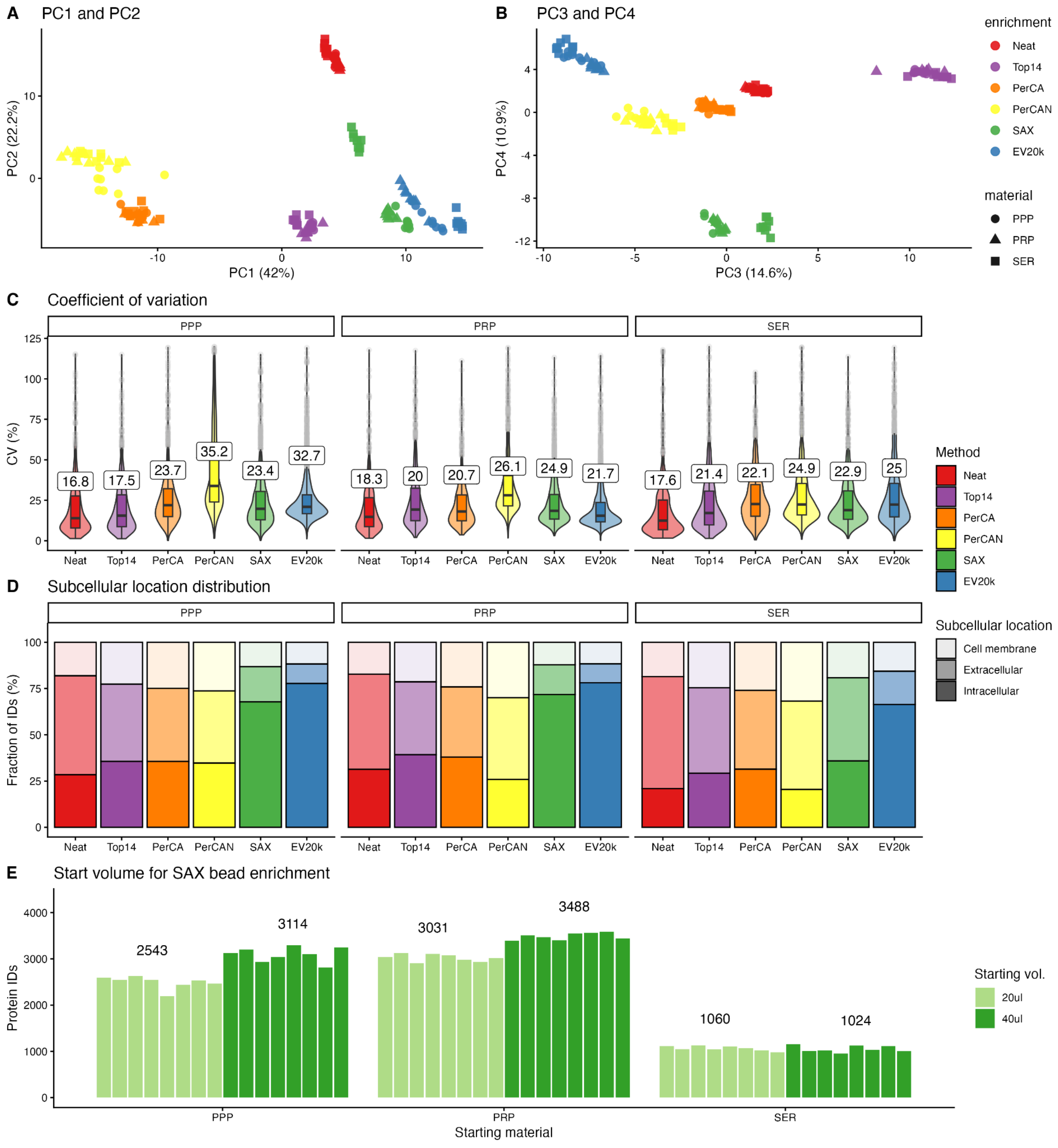

#### Sup. Figure 3:

A: Histogram of donor age in the cohort.

B: Distribution of male (M) and female donors in the cohort.

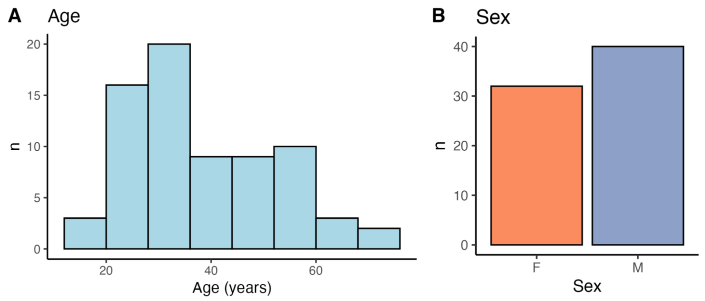


#### Sup. Figure 4

A: All hospital measurements and reference values. The values below LOD is marked as stars. B: Missing values for all hospital (SER) and MS measurements (PRP).

B: Proportion of missing values for each of the 10 proteins compared to clinical routine measurements.

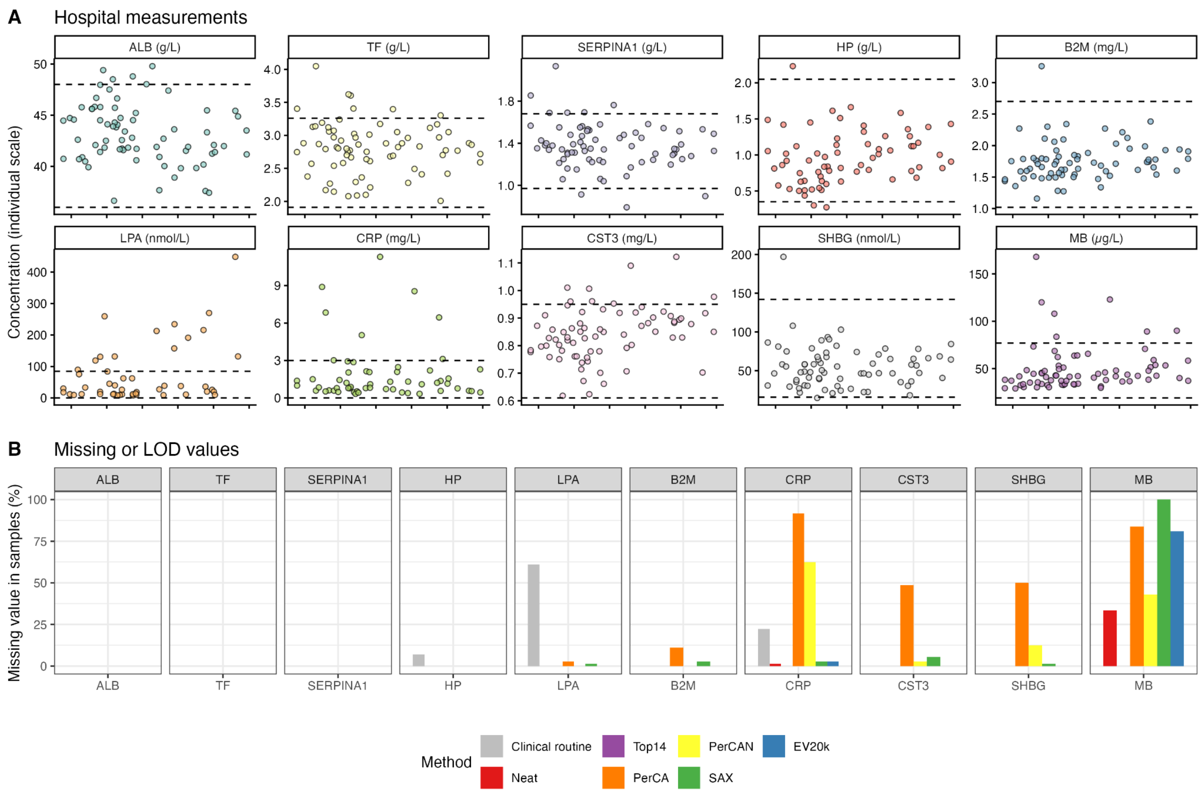

#### Sup. Figure 5

Single protein correlations of clinical routine measurement of MS measurement after each workflow. The Pearson correlation coefficient is shown for each method.

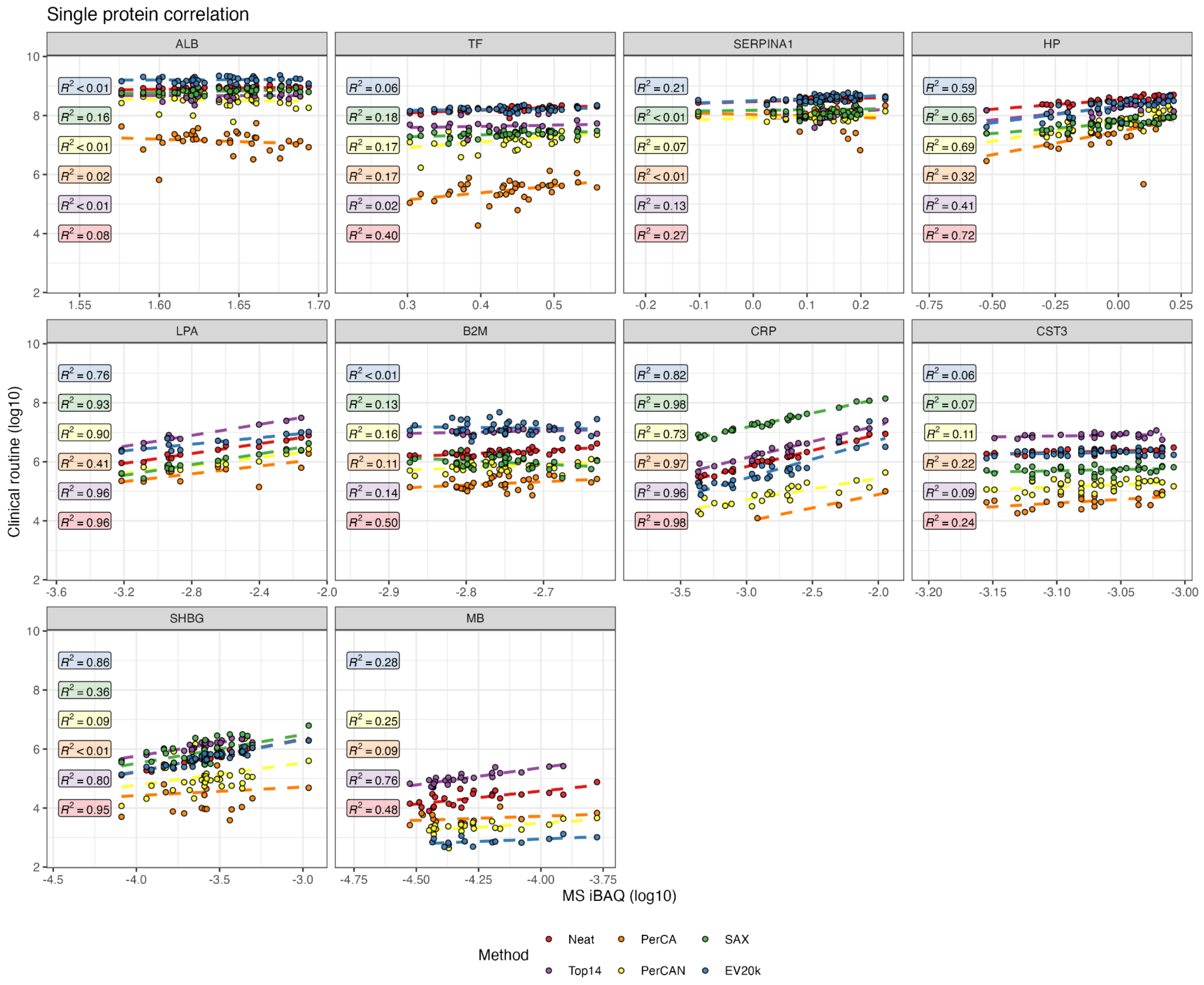

#### Sup. Figure 6

A: Comparisons of protein measurements in male (M) vs female (F) in clinical routine measurements. The p-values are calculated by a student's t test.

B: Comparison of protein measurement vs age. The p values are calculated by fitting a linear regression model.

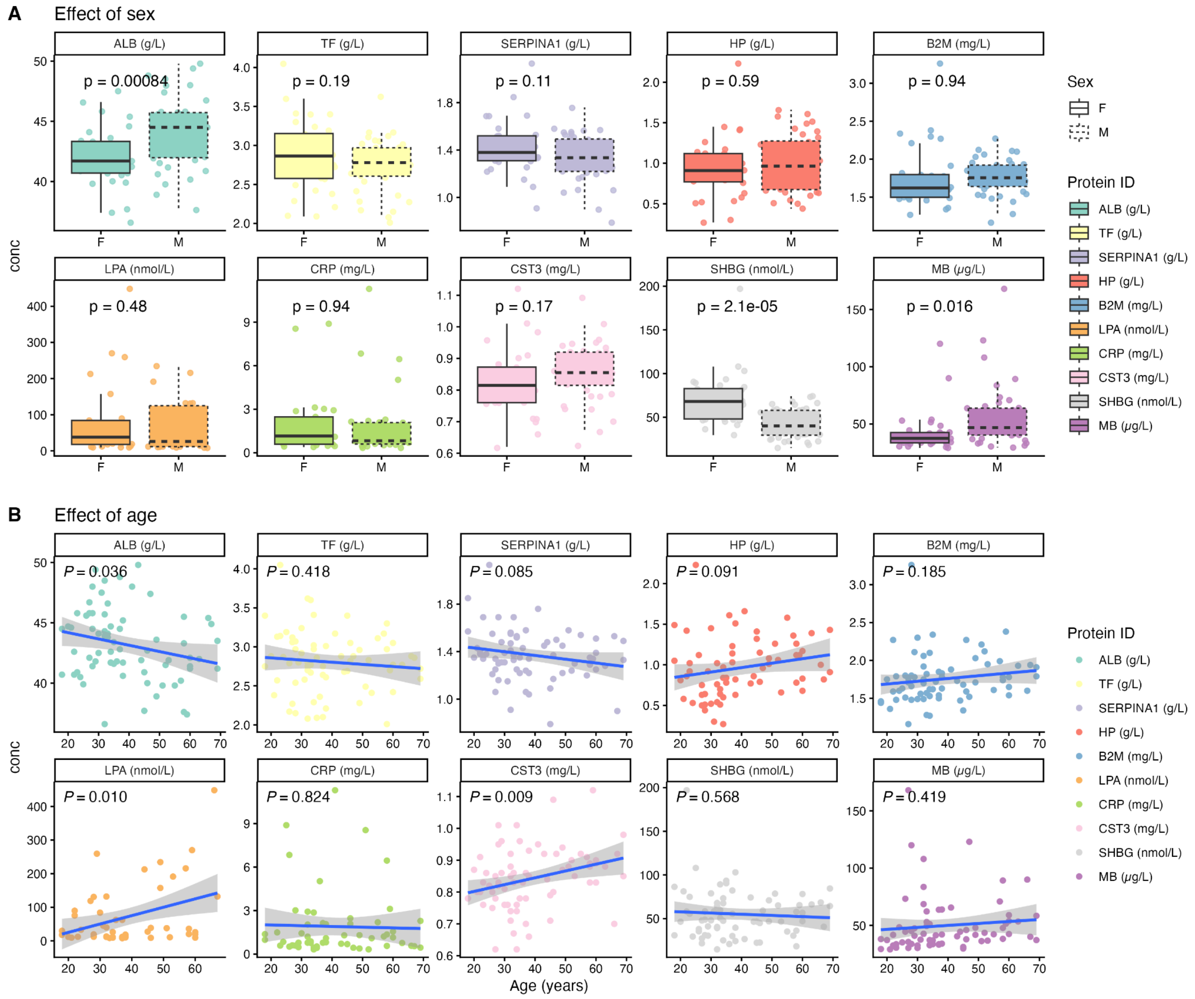
